# A novel rotavirus reverse genetics platform supports flexible insertion of exogenous genes and enables rapid development of a high-throughput neutralization assay

**DOI:** 10.1101/2023.02.15.528690

**Authors:** Jiajie Wei, Scott Radcliffe, Meiqing Lu, Yuan Li, Jason Cassaday, William Newhard, Gwendolyn J Heidecker, William A Rose, Xi He, Daniel Freed, Michael Citron, Amy Espeseth, Dai Wang

## Abstract

Despite the success of rotavirus vaccines, rotaviruses are still one of the leading causes of diarrheal diseases, resulting in significant childhood morbidity and mortality especially in low- and middle-income countries. Reverse genetics system enables the manipulation of the rotavirus genome and opens the possibility of using rotavirus as an expression vector for heterologous proteins such as vaccine antigens and therapeutic payloads. Here, we identified three positions in rotavirus genome, the C terminus of NSP1, NSP3 and NSP5, for reporter gene insertion. By using rotavirus expressing GFP, we developed a high-throughput neutralization assay and revealed the pre-existing immunity against rotavirus in human and other animal species. Our work shows the plasticity of rotavirus genome and establishes a high-throughput assay for interrogating humoral immune responses, benefiting the design of next-generation rotavirus vaccines and the development of rotavirus-based expression platforms.

## Introduction

Although rotavirus vaccines have reduced rotavirus related childhood morbidity and mortality substantially worldwide^1^, rotaviruses are still one of the most common causes of diarrheal diseases in children with a higher disease burden in developing countries. Rotaviruses lead to 128,500-215,000 deaths in children under 5 years old annually^2,3^. The mechanisms underlying rotavirus vaccine induced protection are not fully understood, partially due to limitations in current animal models of rotavirus infection and disease. The observed lower vaccine efficacy in low-income countries has been attributed to multiple factors including higher levels of maternally derived antibodies, different intestinal microbiome resulted from chronic enteropathy, and/or poor nutritional status^4^. To develop the next-generation vaccines with improved safety and efficacy, a better understanding of preexisting immunity including neutralizing antibodies in human and animal models and its impact on vaccine efficacy is needed.

Rotaviruses are double-stranded, segmented RNA viruses. The 11 genome segments encode 12 viral proteins including 6 non-structural proteins (NSP1-NSP6) and 6 structural proteins (VP1-VP4, VP6 and VP7). Each segment encodes one open reading frame (ORF) except segment 11 with NSP5 containing an internal ORF for NSP6. The rotavirus particles are composed of three concentric proteins shells (VP2, VP6, VP7) and a spike protein VP4, which spans the VP6 and VP7 layers and extends out from the particle. Rotaviruses are grouped serologically into distinct serogroups based on VP6 reactivity. Group A rotaviruses (RVA), further classified into serotypes defined by VP7 (G) and VP4 [P], cause a majority of disease in human. RVA strains are also found in animals and infection has been shown to be highly species specific.

Rotavirus research had been hindered by a lack of reverse genetics system where defined viral clones can be generated and engineered. Since the first publication of a plasmid only-based reverse genetics system for simian rotavirus strain SA11^5^, multiple groups have used the system to rescue recombinant rotaviruses with different properties such as simian RRV strain, human CDC-9 strain, a murine-like RV strain^6^, human rotavirus strain Odelia^7^, chimeric strains with SA11 backbone and VP4, VP7, and/or VP6 genes from human clinical samples^8^, and SA11 carrying NSP2 phosphorylation mutation^9^. Heterologous protein expression from rotavirus was explored by replacing part of the NSP1 ORF with foreign genes^5^. Later, it was shown that genome segment 7 could be re-engineered to encode NSP3 fused to a fluorescent protein^10^. Domains of SARS-CoV-2 spike protein were also expressed downstream of NSP3^11^, suggesting rotavirus may serve as a vector for gene delivery. Recently, the concept of using recombinant rotaviruses expressing norovirus capsid proteins as a dual vaccine was established in an inface mouse model^12^. A solid understanding of preexisting immunity in human and animal models would further explore the feasibility of using rotavirus to deliver therapeutic proteins or vaccine antigens.

Here, besides NSP3 site that was published before, we identified two additional genomic locations in NSP1, and NSP5 of simian rotavirus strain SA11 for heterologous protein expressing by reverse genetics. Using a recombinant rotavirus expressing GFP from NSP1 (rSA11-GFP), we established a high-throughput rotavirus microneutralization assay. This assay enabled us to determine pre-existing neutralizing antibodies in human and other animal serum samples. We found that out of 20 human donor samples that were surveyed, 19 had detectable neutralization titers against SA11, a serotype G3 virus. All African green monkeys and rhesus monkeys examined showed pre-existing immunity. Other animal models such as rabbit, mouse, guinea pig and cotton rat have either much lower level or no neutralizing antibodies. By identifying multiple novel sites of rotavirus genome for heterologous gene expression, we demonstrated the plasticity of rotavirus genome and developed a high throughput neutralization assay based on a GFP expressing virus.

## Results

### The identification of three positions in rotavirus genome for heterologous gene expression

We aimed to construct rotaviruses expressing heterologous proteins in addition to the full set of rotavirus proteins. A plasmid only system to rescue simian rotavirus strain SA11 enables the engineering of rotavirus^5^. Because naturally occurring human rotaviruses that contain rearranged genomes can package up to 1,800 additional base pairs in virus particles^13^, and rearranged genome segments 7 and 11 with open reading frame (ORF) sequence repeats at the C terminus of NSP1 and NSP5, respectively, are preferentially packaged into rotavirus^14^, we reasoned that these two positions could tolerate foreign gene insertion.

To systematically explore genomic positions that support heterologous protein expression, we fused a green fluorescent protein (GFP) after a 2A self-cleaving peptide to the C terminus of every nonstructural protein (Fig. 1A) except NSP6 given NSP6 is encoded by an internal ORF inside of NSP5. To keep the potential genome packaging signal at the 3’end of ORF, after the stop codon, 450 bp fragment of NSP ORF 3’ sequence was repeated upstream of the 3’UTR.

**Figure 1.**
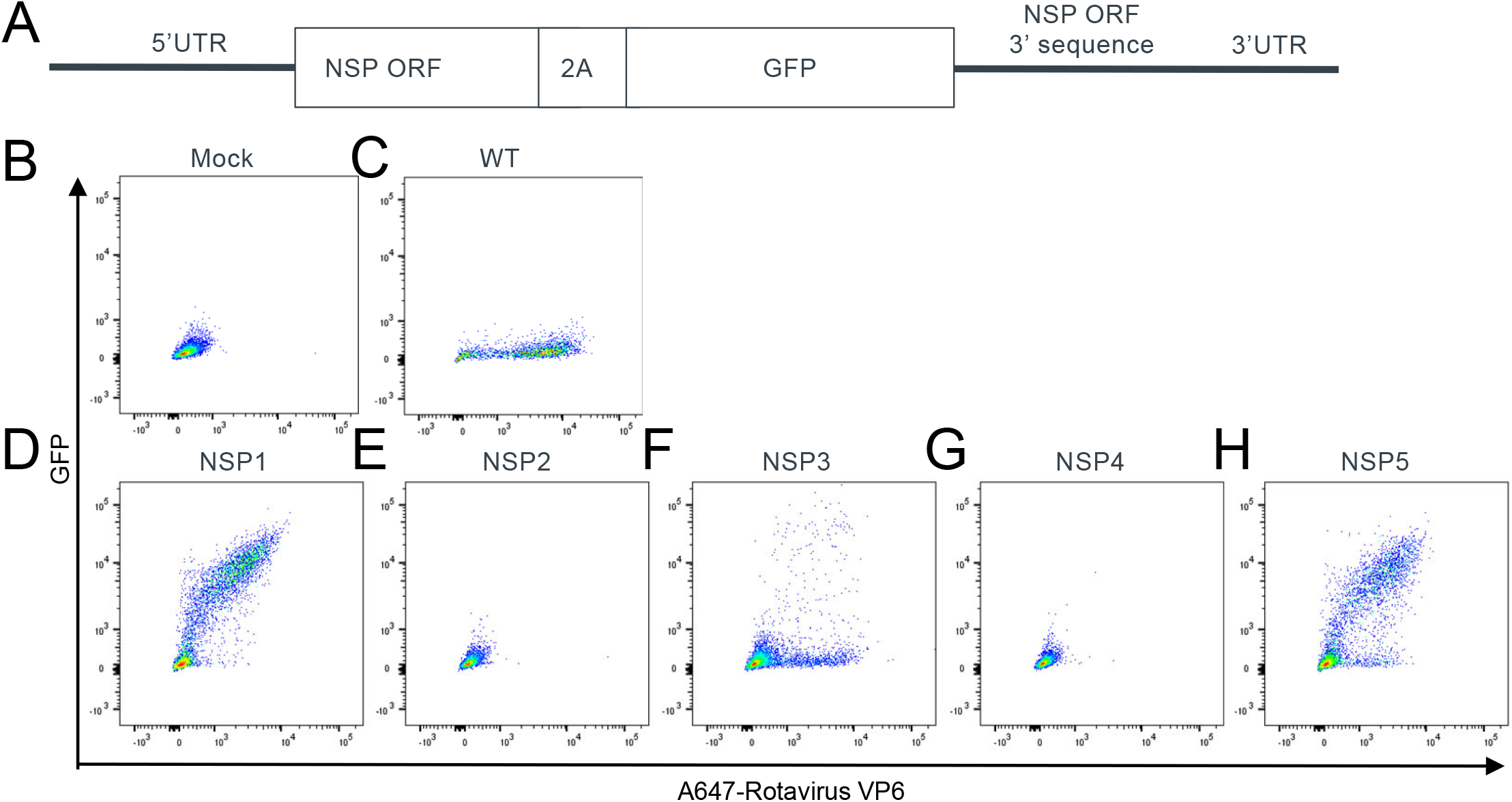
Generation of rSA11 strains expressing GFP from five genome locations. (A) Schematic representation of plasmids used for the rescue of rSA11 viruses encoding GFP. (B-H) Representative flow cytometry analysis of CV-1 cells infected with rSA11 rescue reactions. GFP sequence was inserted at the indicated genome location. Expressions of GFP and rotavirus protein VP6 were examined.

We tested each modified genome segment by replacing the corresponding wild type (WT) SA11 genome segment in the reverse genetics system^5^. We transfected cells with plasmids, used supernatant to infect the CV1 cells, and then determined expression levels of GFP and rotavirus VP6 protein from CV-1 cells by flow cytometry (Fig. 1B-H). As expected, compared to mocked infected cells, CV-1 cells infected with WT virus showed VP6 signal (Fig. 1B and C). We failed to generate recombinant rotaviruses that contained modified NSP2 or NSP4 as VP6 positive cells were not observed after infecting CV-1 cells with corresponding reverse genetics reactions (Fig. 1E and G). In contrast, double positive cell populations that expressed VP6 and GFP were observed when the rescue reactions contained modified NSP1, NSP3 or NSP5, indicating that we successfully generated recombinant rotavirus with GFP insertion at each of these locations (Fig. 1D, F and H). We thus identified three positions for heterologous gene insertion in rotavirus genome. Among NSP1, NSP3 and NSP5, modified NSP1 and NSP5 led to higher percentage of VP6/GFP double positives (Fig. 1D), suggesting higher efficiency of virus rescue.

We then deleted NSP1 and NSP5 ORF 3’ sequence repeat after the stop codon to avoid potential recombination events. We still observed the double positive cell population that expressed GFP and VP6 (Fig. 2), indicating virus packaging does not require the additional repeated sequence. Similar to recombinant rotaviruses rescued in Fig.1, after deleting the repeated sequence, NSP1 position still resulted in higher percentage of P6/GFP double positive population than NSP5 (Fig. 2B). We thus focused on virus containing GFP downstream of NSP1 without 3’ sequence repeat for further characterization and referred it as rSA11-GFP.

**Figure 2.**
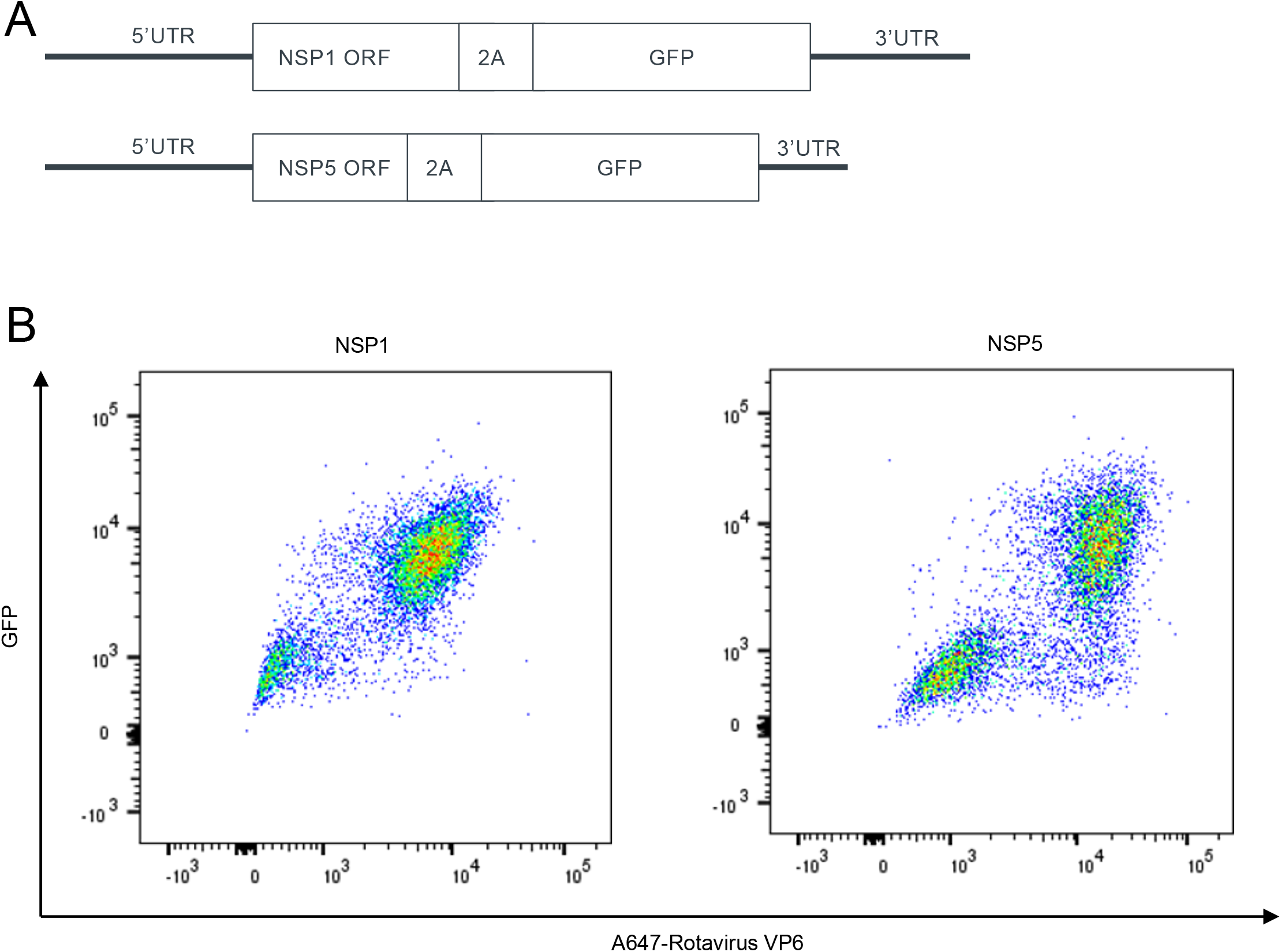
Generation of rSA11 strains expressing GFP downstream of NSP1 or NSP5 without 3’ end ORF repeats. (A) Schematic representation of plasmids used for the recovery of rSA11 virus encoding GFP downstream of NSP1 or NSP5. (B) Representative flow cytometry analysis of CV-1 cells infected with rSA11 strains generated with plasmids shown in (A). Expressions of GFP and rotavirus protein VP6 were examined.

Using flow cytometry to examine VP6 expression, we also established a flow cytometry-based infectivity assay for determining rotavirus titer (Fig. S1). After infection, CV-1 cells were cultured in media supplemented with FBS but not trypsin to prevent multiple rounds of infection. Based on Poisson distribution, the multiple of infection (MOI) can be calculated after determining the percentage of VP6 positive cells. The titer as infectivity unit per ml (IU/ml) can then be calculated based on the number of cells and the amount of viral stock used at infection.

### Characteristics of rSA11-GFP

To examine the genetic stability of rSA11-GFP, we passaged rSA11-GFP and rSA11-WT on MA104 cells ten times, extracted viral RNA from passage one (reverse genetics product) and ten, and performed RT-PCR using primers flanking the insertion site. RT-PCR products were visualized by gel electrophoresis and sequenced by Sanger sequencing. Fragments migrated to expected sizes (Fig. 3A) and sequencing reactions showed that no mutations were generated for 10 passages. Our results thus indicated that rSA11-GFP was genetically stable.

**Figure 3.**
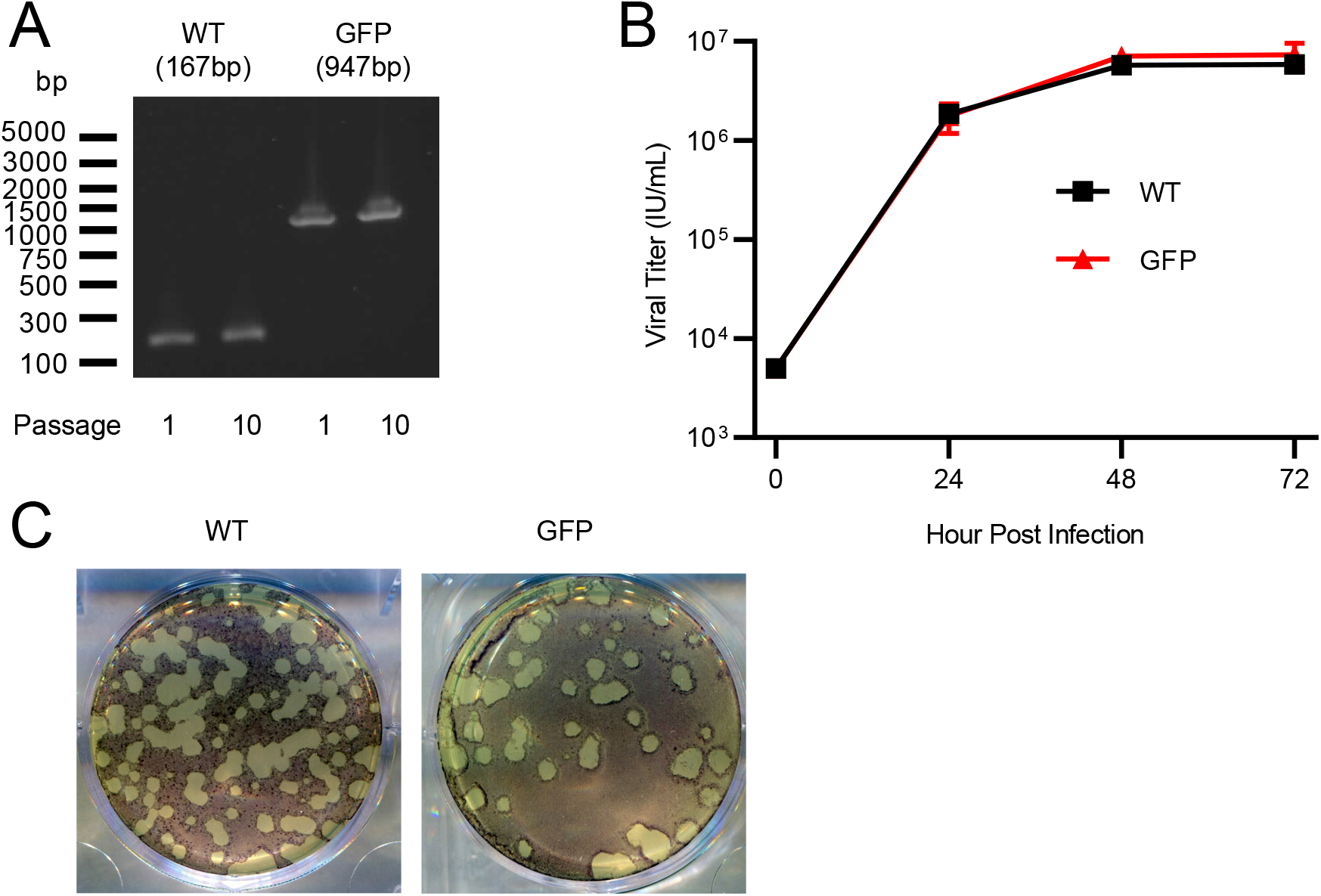
Properties of rSA11-GFP. (A) Genetic stability of rSA11-GFP. rSA11 and rSA11-GFP were serially passaged ten times in MA104 cells and analyzed by RT-PCR using primers flanking the insertion site. The expected band sizes are indicated in parentheses. (B) Growth kinetics of rSA11 and rSA11-GFP. MA104 cells were infected with viruses at an MOI of 0.01 IU/cell and harvested at indicated time points. Virus titer was determined in a flow cytometry-based infectivity assay using CV1 cells. Data are expressed as the mean and range of duplicate samples. (C) Plaque formation on MA104 cells by rSA11and rSA11-GFP. Data representative of three independent experiments.

We also compared the growth kinetics of rSA11-GFP with rSA11-WT. MA104 cells were infected with viruses at an MOI of 0.01 IU/cell and harvested at 24, 48 and 72 h post infection (Fig. 3B). The growth curves of rSA11-GFP and rSA11-WT were indistinguishable, indicating that the insertion of GFP did not affect the fitness of the recombinant virus in vitro. In addition, plaques formed by rSA11-GFP and rSA11-WT were of similar sizes (Fig. 3C), further supporting that the insertion of GFP downstream of NSP1 had no effects on rotavirus replication.

### rSA11-GFP based microneutralization assay

Because traditional neutralization assays such as plaque reduction neutralization test (PRNT) and fluorescent foci reduction neutralization test (FRNT) that relies on antibody staining are time consuming and labor intensive, we sought to develop a microneutralization assay based on GFP signal using rSA11-GFP. In the 96-well plate format, CV-1 cells were cultured overnight before virus infection. rSA11-GFP virus with an MOI of one was mixed with serial diluted animal serum samples for 1 h at 37°C and then the virus/serum mixtures were applied to CV-1 cells for absorption. After overnight incubation, numbers of GFP positive cells were determined to generate neutralization curves. Percentages of inhibition were calculated based on control wells where no animal serum was added. It is known that immunity against rotavirus exists in some monkey colonies naturally^15^. We examined four rhesus monkey serum samples in the 96-well format microneutralization assay and found that, as expected, all four samples neutralized rSA11-GFP with two showing higher neutralizing capacity (Fig. 4A).

**Figure 4.**
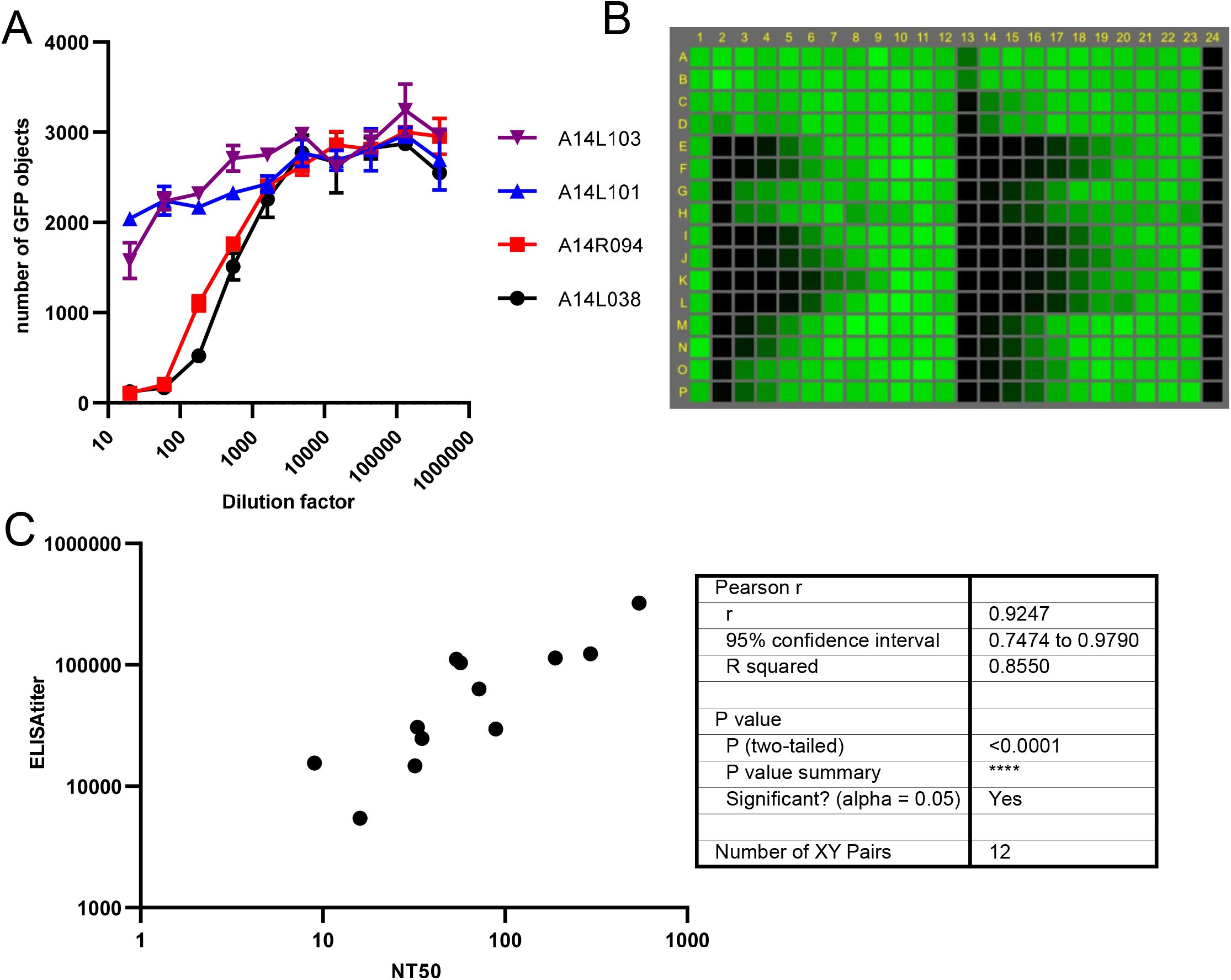
rSA11-GFP based microneutralization assay. (A) Representative serum neutralization curves of four rhesus monkeys. rSA11-GFP, preincubated with serial diluted serum samples, was used to infect CV-1 cells. After overnight incubation, numbers of GFP positive cells were numerated. Median is shown from triplicated wells. (B) The conversion of the neutralization assay to a high-throughput platform. rSA11-GFP, preincubated with serial diluted serum samples, was used to infect CV-1 cells. After overnight incubation a representative review of a 384-well plate was captured. No serum samples were used in the first column and the last column contained mocked infected cells. Wells are colored based on the numbers of GFP positive cells. (C) The correlation of neutralization titers and ELISA titers of serum samples from 12 African green monkeys and its statistical analysis.

We then converted the assay to a higher throughput format by adapting the assay to 384 well plates and eliminating the CV-1 pre-seeding step (Fig. 4B). In the 384-well plate format, CV-1 cells in suspension were applied directly to the virus/serum mixtures for infection. We used this high-throughput assay to examine 12 African green monkeys in our animal facility (Fig. 4C). All animals showed pre-existing antibodies against rotavirus based on neutralization assay and IgG ELISA assay that used SA11 for coating. We observed a wide range of antibody titers with NT_50_ titers ranging from 9 to 545 and ELISA titers ranging from 5,444 to 323,096. Our results indicated that all African green monkeys we examined were seropositive for SA11. It is unlikely that we are detecting maternal antibodies as all monkeys are 2-3 years old. The high level of correlation (r= 0.9247, P<0.0001) between neutralization titer and ELISA titer (Fig. 4C) suggested that either almost all antibodies captured by ELISA were neutralizing antibodies or the proportions of rotavirus antibodies with neutralization capacity were consistent among African green monkeys.

### Pre-existing immunity in human and other animal species

We next determined neutralizing antibodies in human donors by the rSA11-GFP based microneutralization assay (Fig. 5A). Group A rotavirus contains more than 20 VP7 (G) serotypes and more than 10 VP4 [P] serotypes^16^. SA11 was originally isolated from a healthy African green monkey and belongs to G3P5B[2]. Serotypes G1, 2, 3, 4, 9 and 12 are epidemiologically important for human. Although both VP4 and VP7 can elicit neutralizing antibodies, G3 specific antibodies can be revealed by this assay as there is no known P5B[2] human strain. Out of 20 samples we examined, only one did not show neutralization titer above the limit of detection, suggesting the wide prevalence of G3 antibodies in human population. The titers were similar to those of African green monkeys in animal facility and higher than those of 11 rhesus monkeys we examined. It was not a surprise that all rhesus monkeys examined showed neutralization titers as rhesus monkey rotavirus RRV also belongs to G3^16^.

**Figure 5.**
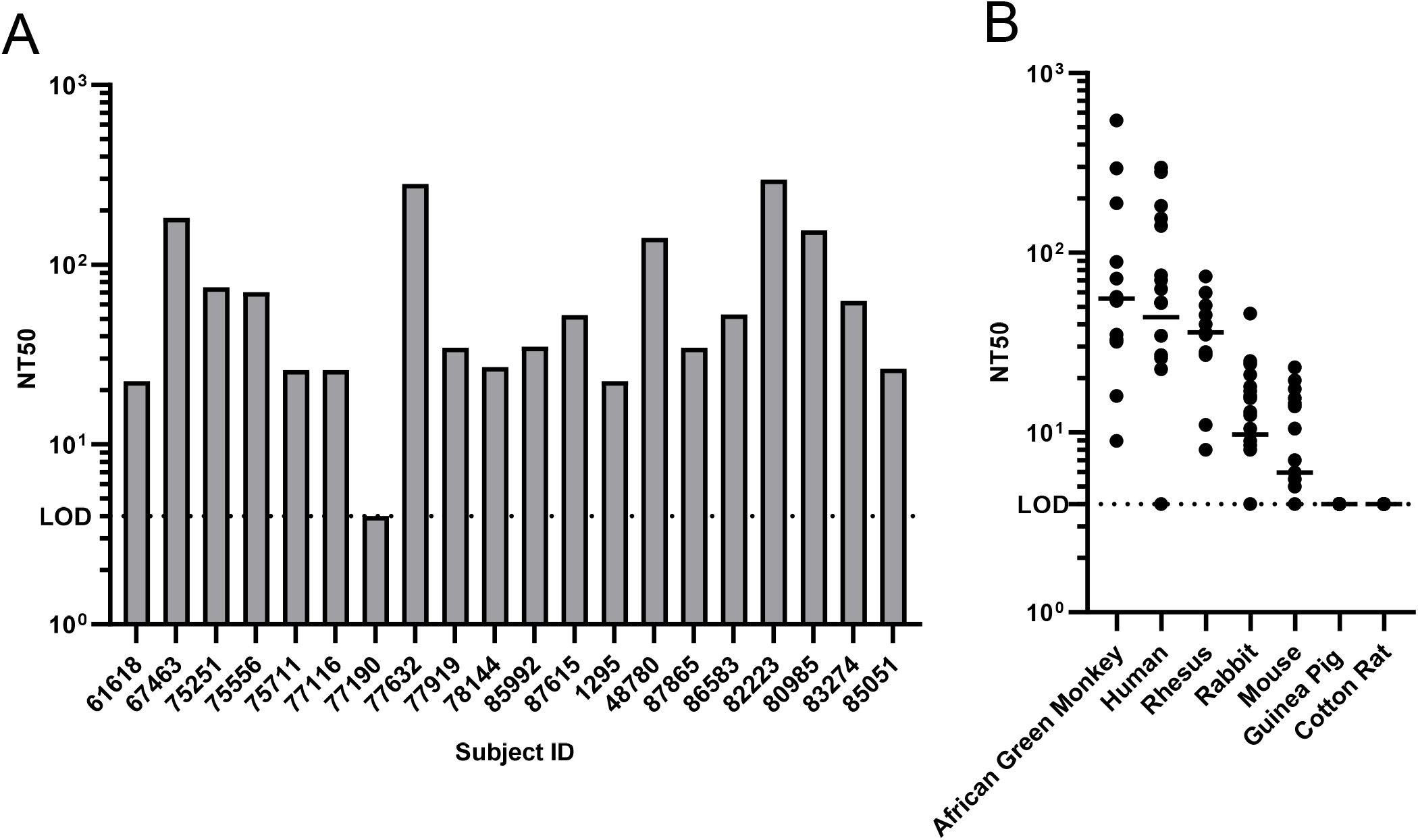
Pre-existing immunity in human and other animal species. (A) Serum neutralization titers of 20 donors. (B) Serum neutralization titers of animal samples from indicated species. The bars indicate the median.

Rotaviruses is a group of viruses impacting a variety of animals, including common animal models used for vaccine or drug development. The microneutralization assay also allowed us to evaluate rabbit, mouse, guinea pig and cotton rat serum samples (Fig. 5B). Several rabbit rotaviruses are G3 serotype viruses and we indeed revealed neutralizing antibodies in many of the rabbits (15 out of 22) although the titers were much lower than those of human or simian. Surprisingly, although there is no known mouse rotavirus strain in the same serogroup as SA11 based on VP4 and VP7, we observed neutralization titers in some of the mouse serum samples (16 out of 25). Guinea pig and cotton rat are being used widely in infectious disease and vaccine research. No rotavirus has been reported in those two species. We did not discover any neutralizing antibodies against SA11 in any guinea pig or cotton rat we examined. In summary, the rSA11-GFP based microneutralization assay enabled us to evaluate pre-existing immunity in several animal species including human in a high-throughput manner.

## Discussion

We extend prior studies^5,7^ on rotavirus reverse genetics systems to identify three positions in monkey rotavirus strain SA11 genome permissive for the insertion and expression of heterologous genes. By using SA11 expressing GFP, we develop a high-throughput microneutralization assay that enables a rapid evaluation of pre-existing antibodies in different animal species including human.

Our results show that rotavirus nonstructural proteins NSP1, NSP3 and NSP5 can tolerate heterologous gene insertion at their C-termini without changing any rotavirus protein coding sequences. Modifying of the C-terminus of NSP1 to express reporter proteins was explored in the original publication on rotavirus reverse genetics, where NSP1 protein was partially deleted to accommodate split GFP fragment GFP11 or NanoLuc ^5^. Our strategy encodes the entire GFP after NSP1 and a 2A cleavage sequence and keeps NSP1 intact as NSP1 impacts intestinal viral replication, pathogenesis, and transmission^17^. In a second publication describing reverse genetics for rotavirus, the C-terminus of NSP3 was fused to heterologous antigens including partial SARS-CoV-2 spike protein ^10,11^. In our experiments, the efficiency of rescuing NSP3-GFP is much lower than that of NSP1-GFP and NSP5-GFP, suggesting that the insertion of foreign sequences at this position renders rotavirus more genetically unstable than at NSP1 or NSP5. The largest insertion at NSP3 has been less than 1.4-kbp. NSP1 and NSP5 positions may tolerate larger insertions, increasing the capability of rotavirus as an expression platform. For the first time, we show that NSP5 can also be manipulated for expressing a heterologous gene.

Neutralizing antibodies are a critical component and indicator of humoral immune response. Traditionally, rotavirus neutralization assay was performed in the format of plaque reduction neutralization test (PRNT) or fluorescent foci reduction neutralization test (FRNT) that relies on immunofluorescence staining. Both assays are laborious and time consuming, impeding an in-depth examination of neutralization antibodies in human and animal models. Microneutralization assays based on GFP or other reporter proteins have been established for a variety of viruses, including human metapneumovirus, human cytomegalovirus, and respiratory syncytial virus ^18-20^. They support high throughput format compatible with robotic system to facilitate basic research and vaccine development.

Rotavirus neutralizing antibodies were discovered against viral proteins VP4, VP7 and VP6 with VP4 and VP7 determining the serogroup and VP6 showing intracellular neutralizing capacity. Here, we developed a microneutralization assay based on rSA11-GFP, a serotype G3P5B[2] virus. The same concept and method can be applied to interrogate other serotypes. Strains including a human clinical isolate have been rescued by reverse genetics ^7^ and chimeric rotaviruses have been generated using SA11 as the backbone ^8^. These results imply that we can either construct rotaviruses from different serotypes expressing GFP or simply generate a panel of rotaviruses containing SA11 NSP1-GFP but VP4, VP7 and VP6 from other strains/serotypes to tease out neutralizing antibodies against each component in animal and human samples.

The high level of correlation between ELISA titers and neutralization titers in our experiments suggest that the microneutralization assay can be used as a surrogate assay for ELISA. By doing so, when interrogating rotavirus humoral immunity, we can bypass the requirement of species-specific secondary antibodies in ELISA and compare neutralizing titers of sera from different animal species. As in previous reports ^15^, all simians we tested showed neutralization titers above the limit of detection. Interestingly, the neutralizing titers of human samples are similar to those of simian with only one negative sample (out of 20), indicating the prevalence of G3 serotype specific antibody in human. Compared to cotton rat and guinea pig where no neutralizing antibodies are detected, more than half mouse samples contain neutralizing antibodies above the limit of detection, although the titers are lower than those of rabbit where several rotaviruses are of G3 type. Our results suggest there might be mouse rotavirus strains with serotypes G3 or P5B[2] that have yet to be discovered.

A high-throughput method to measure neutralizing antibodies enables two types of studies. First, once a panel of recombinant rotaviruses with different serotypes containing NSP1-GFP is established, it provides a tool set for seroepdemiology studies to characterize rotavirus prevalence of different serotypes in human. Second, the method also facilitates the selection of appropriate animal models for preclinical studies. These studies will provide insights into the mechanisms underlying the low efficacies of rotavirus vaccine in developing country to enable a rational design for the next-generation rotavirus vaccine. Importantly, now it is established that rotavirus can express heterologous proteins, these studies will address the potential of using rotavirus as a vector for delivering vaccine antigen or therapeutic payload.

## Materials and Methods

### Cell culture

CV1, MA104, and baby hamster kidney cells expressing T7 RNA polymerase (BHK-T7) were maintained in Dulbecco’s Modified Eagle’s Medium (DMEM) with 10% fetal bovine serum (FBS) and 1% penicillin-streptomycin. All cultures were grown at 37°C in a 5% CO2 incubator.

### Plasmid construction

Sequences of all 16 plasmids used for the generation of wild type SA11 strain were obtained from Addgene (https://www.addgene.org/Takeshi_Kobayashi/)^5^. pUC19 and pV1Jns were the backbones of 11 plasmids each encoding one rotavirus genome segment (pT7/VP1SA11, pT7/VP2SA11, pT7/VP3SA11, pT7/VP4SA11, pT7/VP6SA11, pT7/VP7SA11, pT7/NSP1SA11, pT7/NSP2SA11, pT7/NSP3SA11, pT7/NSP4SA11, and pT7/NSP5SA11) and 5 helper plasmids (pCMV/NSP2, pCMV/NSP5, pCMV/NBVFAST, pCMV/D12L, and pCMV/D1R), respectively. For the expression of GFP, NSP open reading frames were fused with 2A peptide GSGEGRGSLLTCGDVEENPGP and then GFP. After the stop codon, NSP open reading frame sequence repeat was inserted when indicated. All plasmids were synthesized by Genewiz.

### Recombinant rotavirus rescue

Recombinant SA11 (rSA11) strains were generated by reverse genetics as described previously with modifications^5^. Monolayers of BHK-T7 cells in 6-well plates (1 × 10^6^ cells/well) were used for transfection. 16 plasmids (0.75 µg/plasmid except 0.015 µg pCMV/NSVFAST) in 150 µl Opti-MEM were added to 150 µl Opti-MEM containing 12.5 ul Lipofectamine 2000. Transfection complexes were incubated at room temperature for 20 min and then added to BHK-T7 cells drop-wise. 24 h post transfection, culture medium was changed into serum free DMEM. 48 h post transfection, 1.5 × 10^5^ CV1 cells were added to transfected cells and TPCK-treated trypsin (1 mg/ml stock) was added to culture medium so the final concentration was 1 µg/ml. The transfection reaction was monitored daily and harvested when complete cytopathic effect (CPE) was observed (typically three days after the addition of CV1 cells). To generate recombinant viruses with heterologous genes, pT7/NSP1SA11, pT7/NSP2SA11, pT7/NSP3SA11, pT7/NSP4SA11, and pT7/NSP5SA11 were replaced with the corresponding plasmid with heterologous gene insertion. rSA11/GFP was then plaque purified for three rounds on MA104 cells.

### Virus infection

Recombinant viruses were treated with 10 µg/ml TPCK-treated trypsin at 37°C for 1 h. Monolayers of cells were washed with serum free DMEM three times and then infected with trypsin-treated viruses in serum free DMEM at 37°C. After 1 h, inoculums were removed.

### Virus stock preparation

MA104 cells were infected with recombinant viruses and cultured in serum free DMEM containing 1 µg/ml trypsin. After complete CPE was observed, viruses were harvested by three freeze-thaw cycles and filtered through 0.2 micro filters. Amicon Ultra-15 50,000 NMWL centrifugal filter unit was used for virus concentration and purification.

### Plaque assay

MA104 cells were infected with recombinant viruses and overlaid with phenol-red free MEM containing 0.8% agarose and 0.5 µg/ml trypsin. After 4 days, plaques were visualized by adding 5 mg/ml MTT or picked directly for plaque purification.

### Flow cytometry and data analysis

CV1 cells were infected with recombinant viruses and cultured in DMEM supplemented with 10% FBS. After overnight incubation, cells were harvested, fixed with 4% paraformaldehyde, and stained with primary antibody anti-RotaVP6 (UK1, ThermoFisher) and then secondary antibody Alexa Fluor 647 AffiniPure Goat Anti-Mouse IgG (H+L) (Jackson Immuno Research Labs). Staining and washing were performed in Perm/Wash buffer (BD Biosciences). Flow cytometric data were acquired using a BD LSRII flow cytometer (BD Biosciences) and gated on single cells. Data analysis were conducted with FlowJo version 10 software (FlowJo LLC).

### Growth kinetics

MA104 cells were infected with recombinant viruses at a multiplicity of infection (MOI) of 0.01 IU/cell and cultured in serum free DMEM containing 1 µg/ml trypsin. Viruses were harvested at 24, 48, and 72 h post-infection by three freeze-thaw cycles. Virus titer was determined by a flow-cytometry based infectivity assay (Supplementary Fig 1).

### Genetic stability

Viruses were serially passaged on MA104 cells. Monolayers of MA104 cells were infected with viruses and cultured in serum-free DMEM containing 1 µg/ml trypsin. When CPE reached completion, cell culture supernatant was used directly for the next round of infection with 1:1000 final dilution. Viral RNA was extracted from 140 µl supernatant using QIAamp viral RNA kit, and 15 µl RNA was used in the SuperScript IV one-step RA-PCR system with forward primer 5’-CAACGGAGGAACTGATTGAAATGAAGAA -3’ and reverse primer 5’-TTGCCAGCTAGGCGCTACT-3’ following manufacturers’ instructions. PCR reactions were analyzed by 1.2% E-gel (ThermoFisher) along with E-Gel 1 Kb Plus Express DNA Ladder. Sanger sequencing reactions were conducted by Genewiz using primers 5’-GCTACTGATCTCCAACTCAGAAGATG-3’ and 5’-TAGTCTGGACGGTCTTGTGA-3’.

### ELISA

96-well assay plates were coated with SA11 (10^5^ PFU/well) in DMEM at 4°C overnight. The plates were washed once with 300 µL/well Washing Buffer (PBS + 0.05% Tween 20), and then blocked with 200 µL/well Blocking Buffer (Alfa Aesar) at 4°C overnight. Blocked plates were incubated with 100 µL/well a series of 3-fold diluted monkey sera in Blocking Buffer at 4°C overnight. Upon the completion of sera incubation, the plates were washed three times with 300 µL/well Washing Buffer and incubated with 100 µL/well 1:4000 diluted alkaline phosphatase conjugated Goat anti-Rhesus IgG H&L (Southern Tech) in Blocking buffer with 0.1% Tween 20 for 1.5 h at room temperature. After washing the plates three times with Washing Buffer, 100 µL/well Tropix CDP-Star Sapphire II substrate (Applied Biosystem) were added. After incubation at room temperature for 10 min, chemiluminescent signal from each well was read on PHERAstar. The threshold value was 25 times the mean plate background. Interpolated titers were calculated by drawing a line between the last point above the threshold and the first point below the threshold and solving for the fold dilution where that line crosses the threshold.

### rSA11-GFP based micro-neutralization assay

4 × 10^4^ CV1 cells/well were seeded into 96 well plate and cultured overnight. Serum samples were heat inactivated at 56°C for 30 min and serial diluted in PBS. rSA11-GFP (used at MOI = 1) was activated with 10 µg/ml TPCK-treated trypsin at 37°C for 1 h and then mixed with serum at 37°C for 1 h on an orbital shaker. Pre-seeded plates were washed three times with serum-free DMEM, infected with virus/serum mixtures at 37°C for 1 h, and then cultured in phenol red free DMEM supplemented with 10% FBS. For the 384 well neutralization assay, CV1 cells were harvested and washed in serum-free DMEM. 1 × 10^4^ CV1 cells in suspension were added into virus/serum mixtures directly and incubated at 37°C for 1 h. FBS was then added to the plate so the final concentration of FBS is 10%. For both 96 well and 384 well plates, after overnight incubation at 37°C, plates were read by an Acumen HCS reader at 488 nm to determine numbers of GFP positive cells in each well. NT50 was calculated by nonlinear four-parameter curve fitting using Prism 8 (GraphPad).

### Animal and human serum samples

Human serum samples were reported before^21^. Rabbit, mouse, guinea pig and cotton rat serum samples were taken from animals housed in the animal facility in the research laboratories at Merck & Co., Inc., West Point, PA, USA. Simian serum samples were collected from animals at the New Iberia Primate Research Center (NIRC-New Iberia, LA). All studies were conducted in accordance with relevant guidelines using protocols approved by NIRC and the Institutional Animal Care and Use Committee of Merck & Co., Inc., Kenilworth, NJ, USA.

## Figure legends

**Figure 1. Generation of rSA11 strains expressing GFP from five genomic positions**. (A) Schematic representation of plasmids used for the rescue of rSA11 viruses encoding GFP. (B-H) Representative flow cytometry analysis of CV-1 cells infected with rSA11 rescue products. GFP sequence was inserted at the indicated positions. Expression levels of GFP and rotavirus protein VP6 were examined.

**Figure 2. Generation of rSA11 strains expressing GFP at the C terminus of NSP1 or NSP5 without 3’ ORF repeats**. (A) Schematic representation of plasmids used for the recovery of rSA11 viruses encoding GFP downstream of NSP1 or NSP5. (B) Representative flow cytometry analysis of CV-1 cells infected with rSA11 strains generated with plasmids shown in (A). Expression levels of GFP and rotavirus protein VP6 were examined.

**Figure 3. Properties of rSA11-GFP**. (A) Genetic stability of rSA11-GFP. rSA11 and rSA11-GFP were serially passaged ten times on MA104 cells and analyzed by RT-PCR using primers flanking the insertion site. The expected band sizes are indicated in parentheses. (B) Growth kinetics of rSA11 and rSA11-GFP. MA104 cells were infected with viruses at an MOI of 0.01 IU/cell and harvested at indicated time points. Virus titer was determined in a flow cytometry-based infectivity assay using CV1 cells. Data are expressed as the mean and range of duplicates. (C) Plaque formation on MA104 cells by rSA11and rSA11-GFP. Data representative of three independent experiments.

**Figure 4. rSA11-GFP based microneutralization assay**. (A) Representative serum neutralization curves of four rhesus monkeys. rSA11-GFP, preincubated with serial diluted serum samples, was used to infect CV-1 cells. After overnight incubation, numbers of GFP positive cells were numerated. Median is shown from triplicated wells. (B) The conversion of the neutralization assay to a high-throughput platform. rSA11-GFP, preincubated with serial diluted serum samples, was used to infect CV-1 cells. After overnight incubation a representative review of a 384-well plate was captured. No serum samples were used in the first column and the last column contained mocked infected cells. Wells are colored based on the numbers of GFP positive cells. (C) The correlation of neutralization titers and ELISA titers of serum samples from 12 African green monkeys and its statistical analysis.

**Figure 5. Pre-existing immunity in human and other animal species**. (A) Serum neutralization titers of 20 human donors. (B) Serum neutralization titers of animal samples from indicated species. The bars indicate the median.

**Supplementary Figure 1.**
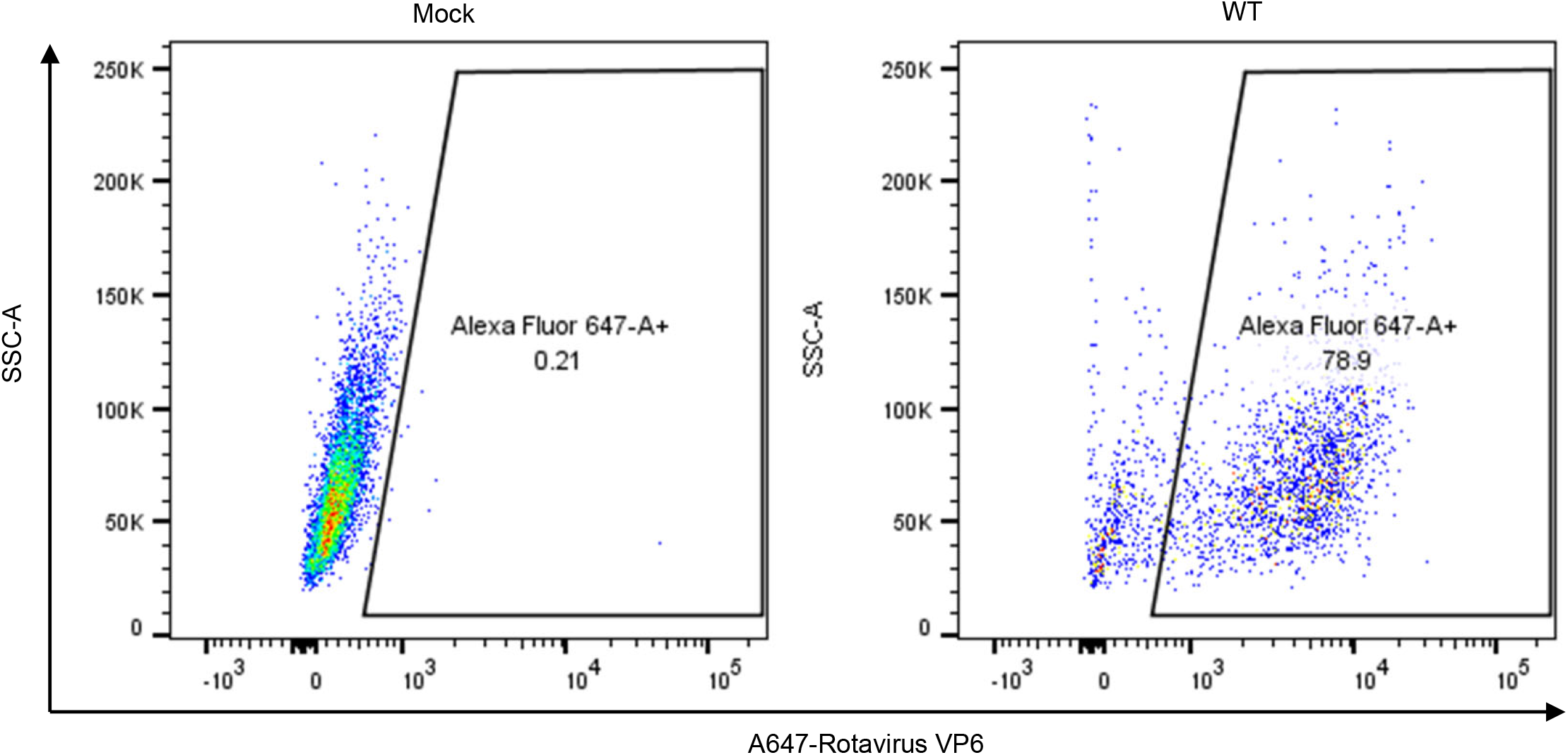
Flow cytometry-based infectivity assay. Mock infected cells (left) and rSA11 infected cells (right) were stained with rotavirus VP6 antibody. MOI was calculated using the percentage of VP6 positive cell population base on Poisson distribution. IU/ml was calculated by IU/ml = (# of cells at infection) × [MOI / (ml of viral stock used at infection)]

## Notes

**Funding Source** This work was supported by Merck Sharp & Dohme Corp., a subsidiary of Merck & Co., Inc., Kenilworth, NJ, USA.

**Declaration of Potential Conflicts of Interest:** All authors are employees of Merck Sharp & Dohme Corp., a subsidiary of Merck & Co., Inc., Kenilworth, NJ, USA. A provisional patent application on the discoveries of this work has been filed.

### Competing Interest Statement

The authors have declared no competing interest.

